# Multivoxel Pattern Analysis Reveals Dissociations Between Subjective Fear and Its Physiological Correlates

**DOI:** 10.1101/515973

**Authors:** Vincent Taschereau-Dumouchel, Mitsuo Kawato, Hakwan Lau

## Abstract

In studies of anxiety and other affective disorders, objectively measured physiological responses have commonly been used as a proxy for measuring subjective experiences associated with pathology. However, this commonly adopted ‘biosignal’ approach has recently been called into question on the grounds that subjective experiences and objective physiological responses may dissociate. We performed machine-learning based analysis on functional magnetic resonance imaging (fMRI) data to assess this issue in the case of fear. Participants were presented with pictures of commonly feared animals in an fMRI experiment. Multivoxel brain activity decoders were trained to predict participants’ subjective fear ratings and their skin conductance reactivity, respectively. While subjective fear and objective physiological responses were correlated in general, the respective whole-brain multivoxel decoders for the two measures were not identical. Some key brain regions such as the amygdala and insula appear to be primarily involved in the prediction of physiological reactivity, while some regions previously associated with metacognition and conscious perception, including some areas in the prefrontal cortex, appear to be primarily predictive of the subjective experience of fear. The present findings are in support of the recent call for caution in assuming a one-to-one mapping between subjective sufferings and their putative biosignals, despite the clear advantages in the latter’s being objectively and continuously measurable in physiological terms.

---

Physiological markers have been used as proxies for psychological states in multiple mental health domains. However, evidence accumulated over the years called into question the relationship between some subjective mental states and their proposed physiological markers. For example, in the case of pain, it is well established that subjective nociceptive experiences can occur without any obvious peripheral physiological manifestations ^1^. As a result, the self-reported subjective experience remains to this day the gold-standard in pain assessment ^2^.

Currently, a similar debate is taking place concerning fear and anxiety ^3–5^. In that literature, physiological reactivity to threat has been considered a reliable objective proxy for the subjective experience of fear ^6^. The reliance on such physiological measures proved to be quite successful and they are now included in numerous studies on fear and anxiety ^7^. Specifically, the neural network involved in physiological reactivity is currently one of the primary neurobiological targets for the pharmacological treatment of anxiety disorders ^8^.

However, some authors suggest that physiological reactivity (as commonly indexed by skin conductance and amygdala reactivity) might represent automatic, defensive responses that aren’t necessarily conscious ^3,4,9^. On this view, studying these physiological defensive responses may not cover all the relevant mechanisms involved in the subjective suffering that is central to fear and anxiety disorders. Accordingly it is argued that, an overemphasis on objective physiological biosignals might slow down the development of new therapeutic options ^4^. This position remains controversial as others have pointed out that multiple lines of evidence actually indicate a high correlation between subjective fear reports and physiological responses, notably in the amygdala ^5^. Here, we attempt to bring in evidences to arbitrate this debate using human functional neuroimaging.

Specifically, our goal is to study if the brain representation of subjective fear ratings dissociates from the representation of objective physiological reactivity (i.e. skin conductance response to feared images). To do so, we focused on naturally occurring instead of conditioned fears. One advantage is that these representations are likely to reflect more closely the brain mechanisms involved in anxiety disorders such as naturally occuring phobia. We constructed a functional magnetic resonance imaging (fMRI) experiment to present as many as 3,600 images of the most commonly feared animals, some neutral animals, as well as some man-made objects as controls (see Fig. 1).

**Figure 1.**
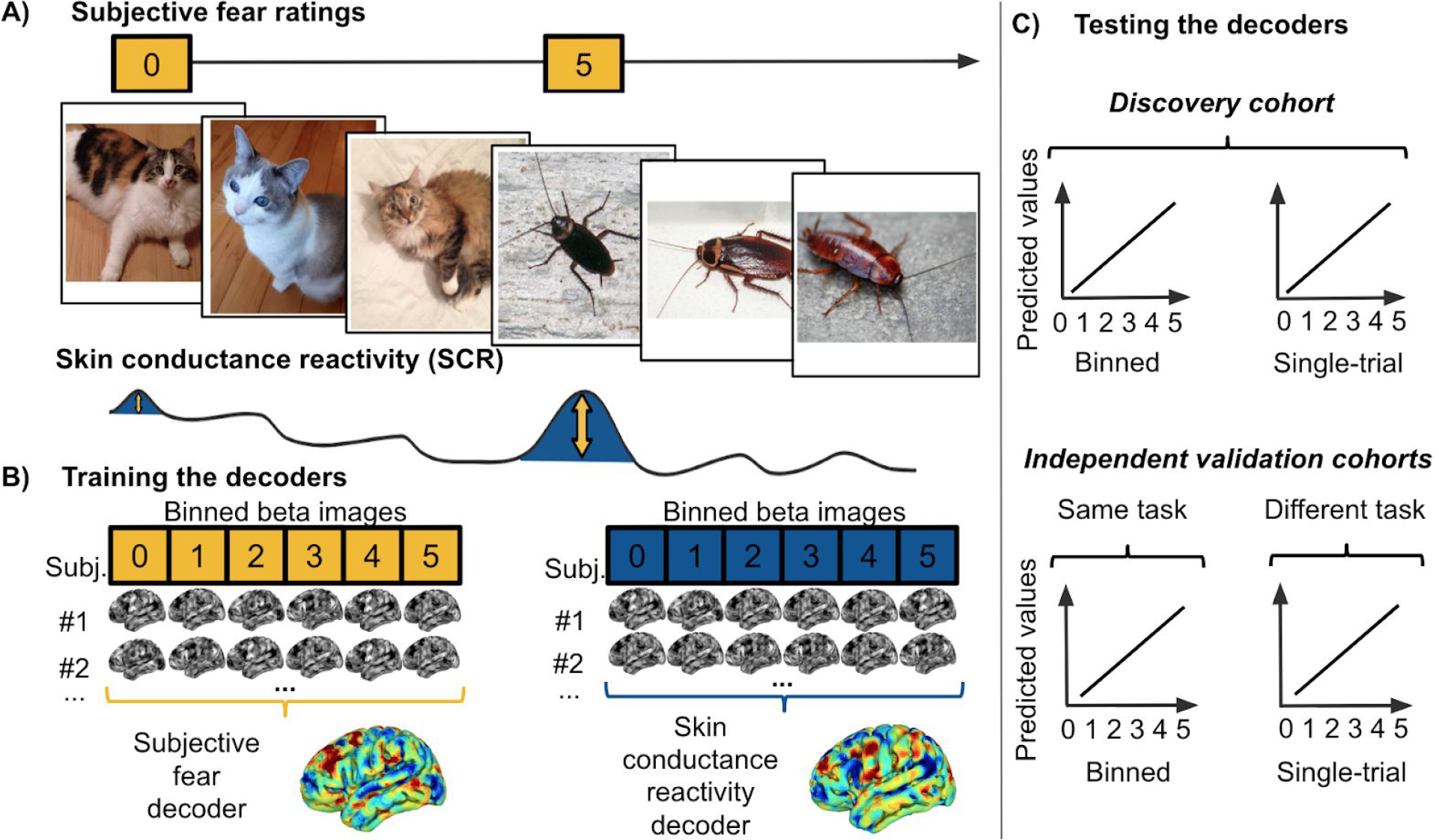
Experimental design and decoding procedure. A) An example sequence of events in the fMRI procedure. We recorded functional brain activity and electrodermal activity during the presentation of images depicting 30 animal categories as well as men-made objects. Akin to the clinical assessment of fear, subjective fear ratings of each of the 30 animal categories were collected outside of the fMRI scanner using a 6-point Likert scale. We estimated brain responses to the first picture of each chunks of a given animal category. B) The estimated brain responses were averaged according to their categorical fear ratings (left) and their skin conductance reactivity (right). This process created binned beta images representing the levels of each outcome for each participant. The binned beta images of the discovery cohort were used to train the decoders. The unthresholded weight maps of the whole-brain decoders are displayed. C) The performance of the decoders were tested in the discovery cohort (both on binned and single-trial data) as well as in independent validation cohorts not included in the training of the decoder. This procedure allowed to estimate the generalization of the decoders to new datasets. The first independent cohort included new participants (N = 12) performing the same task as the one performed by the discovery cohort. The second independent cohort (N = 17) performed a different experimental task where pictures of feared animals were also presented (see Supplementary Methods and Results)

We used a machine learning approach ^10–12^ to train multivoxel brain decoders to predict either objective physiological reactivity or subjective fear reports. To do so, we leveraged whole-brain data in order to determine the patterns of voxel activities that are the most predictive of each outcome (i.e., levels of fear and levels of skin conductance reactivity). The accuracy of these decoders was tested using leave-one-subject-out cross-validation as well as 2 independent validation datasets (N = 12 and N = 17) (see Fig. 1 c). We also aimed at determining if some brain regions are preferentially involved in the prediction of either the subjective or physiological measures. As such, we established where in the brain it was possible to predict one outcome with a better accuracy than the other. This was achieved by comparing the predictions of both decoders within brain regions.

To anticipate, we found that the representations of subjective fear and skin conductance reactivity present some overlap but also some differences in the brain. Specifically, regions previously associated with defensive responses (such as the amygdala) present a preference in the prediction of defensive responses while some higher order frontal regions, previously associated with conscious perception and metacognition, appear primarily involved in the prediction of the subjective fear reports.

In our study, we aimed to estimate the brain activity associated with varying levels of subjective fear and physiological reactivity. To do so, we presented a reasonably large number of images from a broad range of animal categories grouped in chunks of 2, 3, 4 or 6 images of the same category. Only the first image of each chunks were modeled in the fMRI analyses because these images could be attributed both a subjective fear rating and a level of skin conductance reactivity (see Methods). The subjective fear ratings were established before the fMRI procedure. Participants were asked to use a 6-point Likert scale to determine their fear of each of the 30 animal categories. This assessment was achieved without presenting any fearful stimuli and is similar to typical approaches used in clinical settings. Skin conductance reactivity was established during the fMRI session using standard analytical procedures (see Methods).

As expected based on previous literature, subjective fear ratings and skin conductance reactivity were correlated (r(28)= .43; *P* = .02; 95% CI: 0.08-0.69; R^2^ = 0.19; two-sided) (see Fig. 2 and Methods). Both outcomes also presented some level of variability. At the group level, 59% of trials were associated with a certain level of fear (4.6% very high fear, 11.0% high fear, 12.8% moderate fear, 15.9% low fear, and 14.7% very low fear) while 41% were associated with no fear. Regarding skin conductance reactivity, 28.21% of trials were considered to present a certain level of reactivity (> .2 microsiemens) while 71.79% did not (see supplementary Methods).

**Figure 2.**
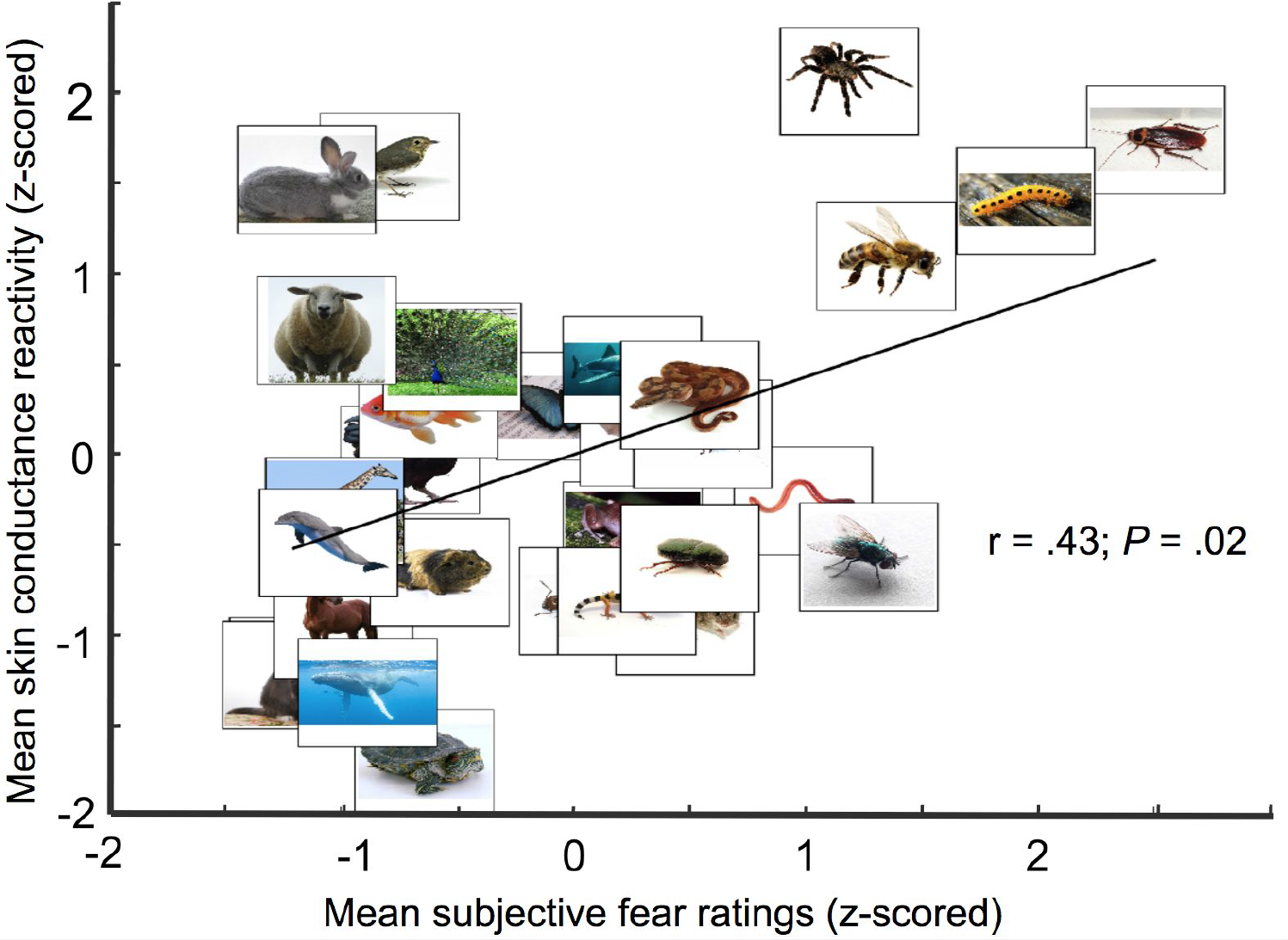
Skin conductance reactivity is correlated with the subjective fear ratings. Within each category, subjective fear ratings and mean skin conductance reactivity were averaged at the group level and standardized (see Methods). As expected, skin conductance reactivity was correlated with subjective fear ratings (r(28)= .43; *P* = .02; 95% CI: 0.08-0.69; R^2^ = 0.19; two-sided).

To train whole-brain decoders, we created 2 datasets by binning (i.e., averaging) together within-subject beta images either as a function of individual fear ratings (0 = “No Fear” to 5 = “Very High Fear”) or as a function of skin conductance reactivity (according to individual quintiles of the reactivity) (see Fig. 1 and Methods). This procedure allowed both to remove the effect of outliers and to capture the within-subject variability of each measure.

In a cross-validation procedure, we trained a support vector regression decoder on the data of N-1 participants and tested the accuracy of the decoder to predict the left-out participant (i.e., leave-one-subject-out cross-validation approach). This procedure was achieved iteratively in order to obtain predicted values for all participants. We established both the sensitivity (e.g., can we predict accurately the subjective ratings of fear?) and the specificity (e.g., can we predict the subjective ratings with the skin conductance reactivity decoder?) of each whole-brain decoders. The sensitivity was established using the area under the ROC curve (AUC) of the predicted values (see Methods). The specificity was determined by testing each decoder using the dataset of the other outcome (e.g., testing the subjective fear rating decoder using the skin conductance dataset). This process, which we call “cross decoding”, can reveal similarities between brain representations if the results reveal above-chance performances. We also determined the performance of the decoders trained with binned beta images in the prediction of single-trial (i.e., unaveraged) beta images. This was achieved also using aleave-one-subject-out cross-validation procedure (i.e., training with binned beta images and testing with single-trial data of the left-out participant).

Figure 3a shows the discrimination accuracy of the subjective fear rating (Fig.3a left panel) and skin conductance reactivity decoders (Fig.3a, right panel). Both decoders present a high level of sensitivity in the prediction of binned images (AUCs ~ .85) as well as a good sensitivity (AUCs ~.62) in the prediction of single-trial images (See Fig. S1 and Supplementary Methods and Results). Both decoders also showed some cross decoding capacity as indicated by above-chance classification of the binned images (dashed lines in Fig. 3 correspond to p = .05).

**Figure 3.**
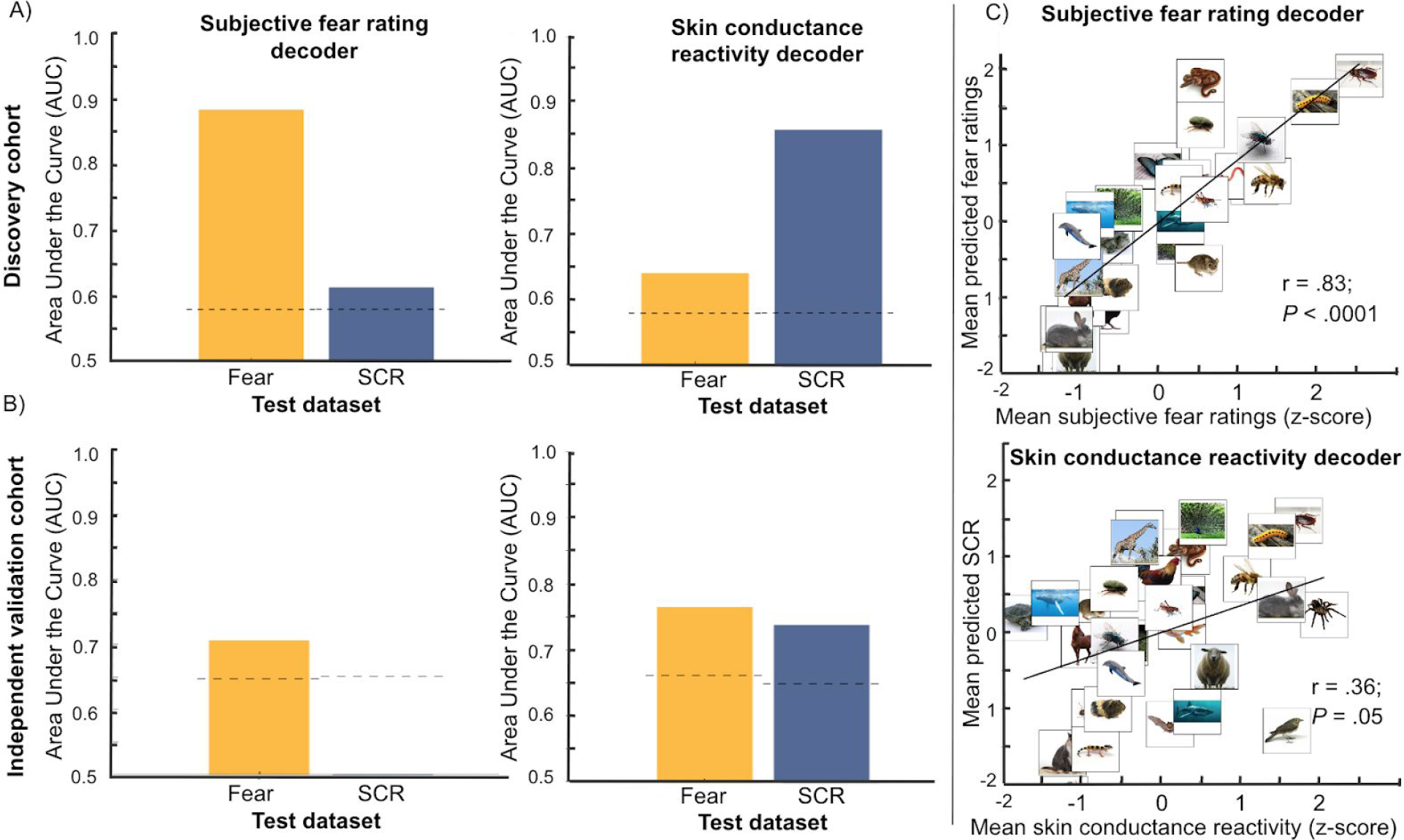
Whole-brain decoders of subjective fear and skin conductance reactivity. A) Both whole-brain decoders presented a good sensitivity when tested on the dataset they were trained to predict (e.g., subjective fear decoder predicting the fear dataset). The cross-decoding procedure (e.g., predicting skin conductance reactivity using the subjective fear decoder) also revealed that both decoders can generalize to some extent to the other dataset. Dashed lines represent the critical value (p = .05) determined using a permutation test. B) Both whole-brain-decoders also generalized to new data as evidenced by their good capacity to predict the data of the independent validation cohort. Also, the cross-decoding procedure indicated that the skin conductance decoder (right panel) could also predict accurately the subjective fear rating dataset. This was not observed for the subjective fear decoder (left panel). C) The whole-brain decoders were also tested on the categorical beta images of each participant. The predicted values of both decoders correlated with the real values of the outcome.

While leave-one-subject-out cross-validation is a common practice in machine learning, this approach may not reflect the true generalization capability of the decoders ^13^. As such, we estimated the generalization of the decoders using 2 independent validation datasets.

The first dataset included a group of participants (N=12) that went through the same fMRI procedure (i.e., same task) but were not included in the training of the decoder (nested cross-validation ^14^). Figure 3b shows the discrimination accuracy of the subjective fear (Fig.3a left panel) and skin conductance reactivity decoders (Fig.3a, right panel) in the prediction of this independent validation cohort. Both decoders still presented sensitivity (AUCs ~ .70) in the prediction of the outcome they were trained to predict. There was also above-chance classification of the subjective fear dataset using the skin conductance decoder (dashed lines in Fig. 3 correspond to p = .05).

The second independent validation dataset was composed of a subsample of participants from the discovery cohort (N=17) that took part in a new fMRI procedure (i.e., different task) (see Supplementary Methods). In this experiment, participants were asked to assess online their subjective fear of images of feared and non-feared animals. Their skin conductance reactivity was being recorded during this fMRI procedure. We processed single-trial beta images from this experiment and submitted these images to the whole-brain decoders (Supplemental Fig. S2 and Methods). This procedure revealed that both decoders presented weak but statistically significant prediction of the independent validation dataset (See Fig. S2 and Supplementary Methods and Results).

Since participants presented various levels of fear and skin conductance response to each category, the prediction of categorical beta images (e.g., averaged beta images of snakes) by each decoder should also follow the subjective ratings and skin conductance reactivity. To test this hypothesis, we computed mean beta images for each animal category (see Methods) and submitted these beta images to the decoders.

The results indicated that whole-brain decoders presented significant predictions of the categorical beta images. More precisely, at the group level, the predictions of the subjective fear decoder were correlated with the subjective fear ratings (r(28) = .82; *P* < .0001; 95% CI: 0.65-0.91; R^2^ = 0.67; two-sided) (see Fig. 3c, top panel) and the predictions of the skin conductance decoder were correlated with the average skin conductance reactivity of the categories (r(28) = .36; *P* = .05; 95% CI: −0.006-0.63; R^2^ = 0.13; two-sided) (see Fig. 3c, bottom panel).

Another important question pertains to the generalization of the brain decoders to a clinical population. To provide some information regarding this generalization, we included in the study three patients (N = 3) diagnosed with specific phobia of one of the 30 animals (see Supplementary Methods). The performance of the decoders were similar to those expected from the rest of the discovery cohort (see Supplementary Methods and Results).

Taken together, these results indicate that it is possible to develop sensitive whole-brain decoders of subjective fear and skin conductance reactivity. Importantly, our results suggest that both decoders can generalize to some extent to 2 independent validation cohorts as well as to patients diagnosed with specific phobia. Furthermore, the predictions of the decoders appear to correspond to the individual variability in the data as assessed by the prediction of the categorical beta images of each animal categories. While these decoders present some similarities, they also appear to be independent from one another as indicated by the results of the cross-decoding procedure.

Next, we aimed to determine the brain regions differentially involved in the prediction of subjective fear ratings and skin conductance reactivity. Thus, we used the same leave-one-subject-out cross-validation but within predefined brain regions. For this purpose, we used a parcellation of the cortex based on functional connectivity ^15^. We selected the 210 cortical regions of this brain atlas as well as the amygdala and hippocampus, for a total of 214 regions. We compared the performance of the decoder within each region. These statistical comparisons were corrected to account for multiple comparisons (see Methods).

Figure 4 and Table 1 indicate in which regions the predictions of the decoders were statistically different. Interestingly, the significant regions of the middle frontal gyrus (Inferior frontal junction, A8vl, A6vl, A10l) all involved a better prediction of the subjective fear ratings than the skin conductance reactivity (see Fig. 4b, left panel). Other regions presenting such a preference for the prediction of the subjective ratings include the medial superior frontal gyrus, the lateral orbitofrontal gyrus, the inferior temporal gyrus, the fusiform gyrus, the parahippocampal gyrus, the superior parietal lobule, the inferior parietal lobule, the precuneus, and the occipital lobe (see Table 1). Furthermore, other regions such as the amygdala, the insula, and the ventral medial prefrontal cortex appear to be primarily associated with the skin conductance response, while being marginally involved in the prediction of the subjective fear ratings (see Fig. 4b, right panel). Other regions presenting such a preference are the lateral inferior frontal gyrus, the superior parietal lobule, the paracentral lobule and the postcentral gyrus. These results suggest that the subjective experience of fear might involve brain processes partly distinct from those involved in the production of the skin conductance response.

**Figure 4.**
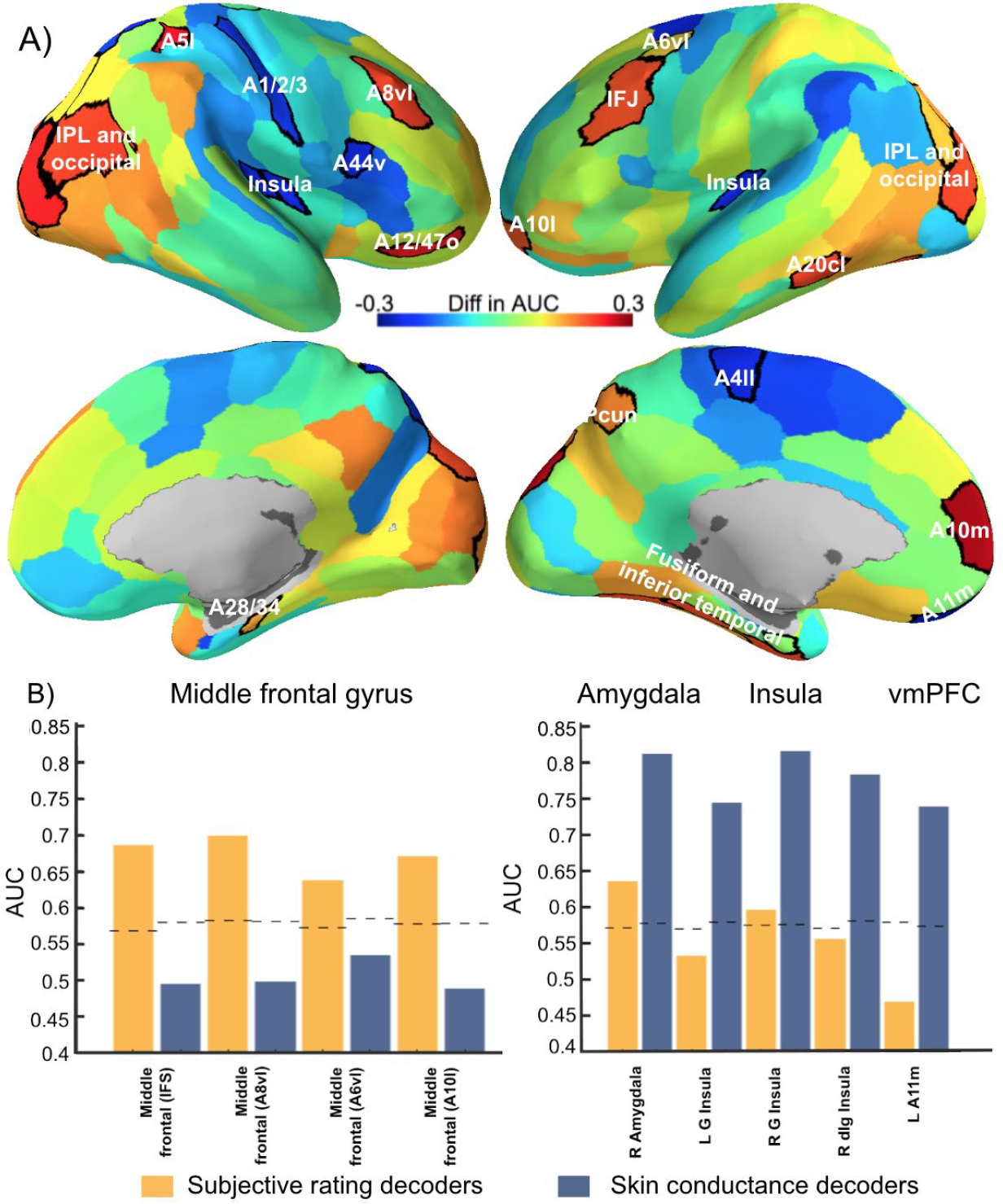
Brain regions presenting a significant difference in the prediction of the subjective ratings and skin conductance reactivity. A) A positive difference in the area under the curve indicates a better prediction of the subjective ratings (red-orange regions) while a negative difference indicates a better prediction of skin conductance reactivity (blue regions). The significant regions (p < .05; FDR-corrected) are surrounded by black borders and are listed in Table 1. Brain images were generated using pySurfer (https://pysurfer.github.io/) B) Significant regions of the middle frontal gyrus, amygdala, insula, and ventral medial prefrontal cortex (vmPFC). Dashed lines represent the critical value (p = .05) determined using a permutation test.

**Table 1.**
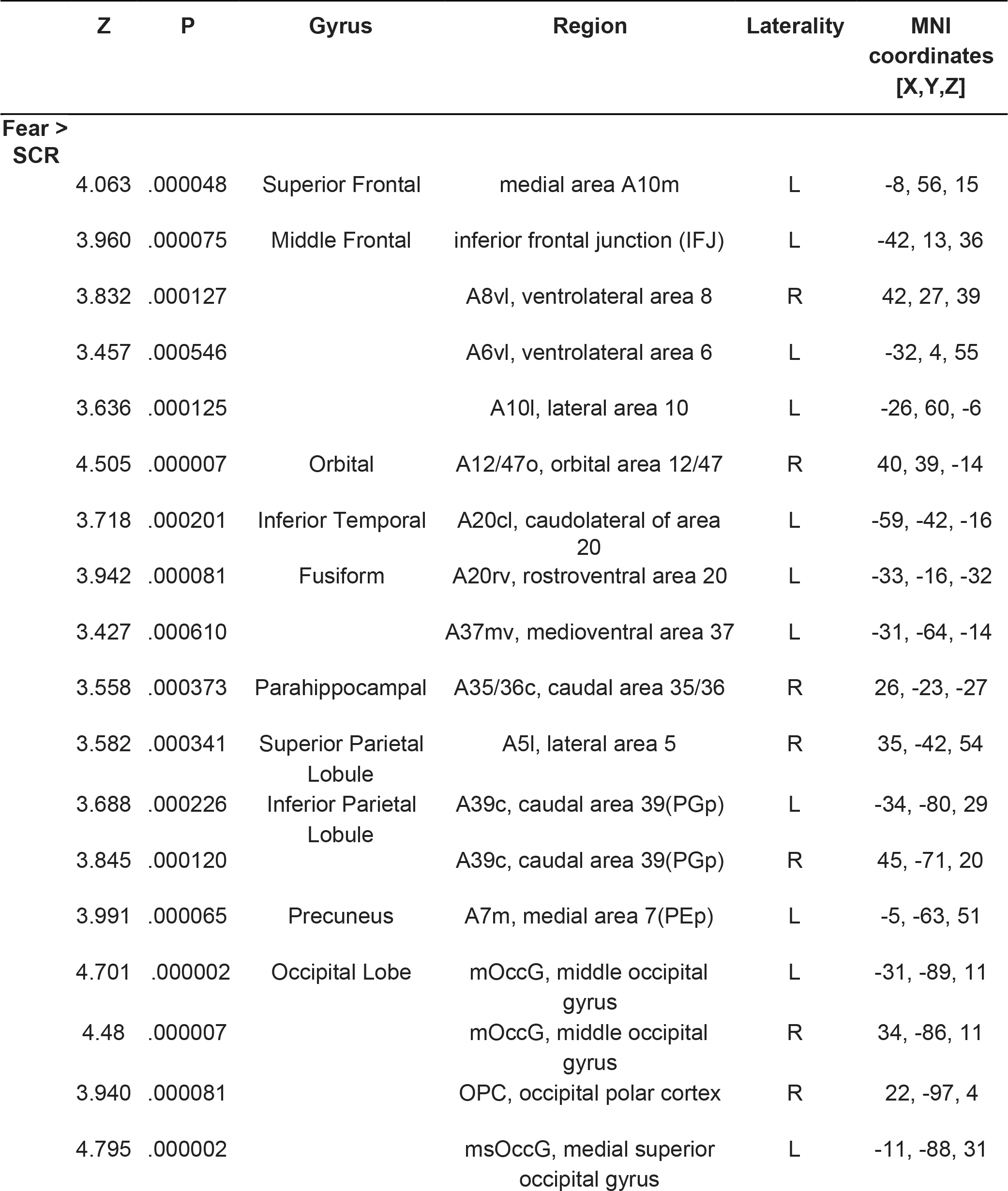
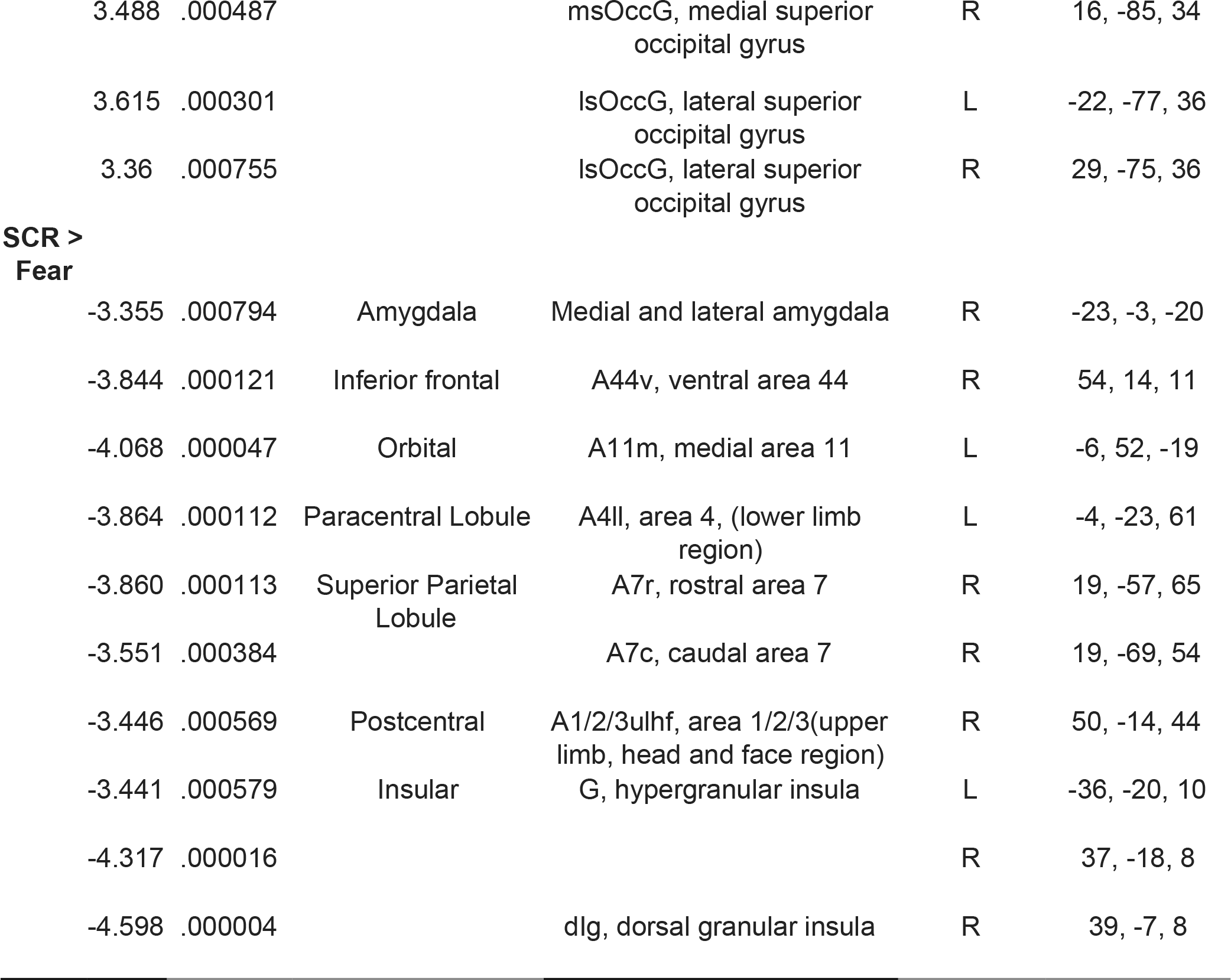
Regions presenting a significant difference in the prediction of subjective ratings and skin conductance responses. Following Fisher’s method ^51^, the Z value can be used to compare directly the correlations between the predicted values of each decoders and the real values.

| Z | P | Gyrus | Region | Laterality | MNI coordinates<br>[X,Y,Z] |
| --- | --- | --- | --- | --- | --- |
| **Fear >**<br>**SCR** |  |  |  |  |  |
| 4.063 | .000048 | Superior Frontal | medial area A10m | L | -8, 56, 15 |
| 3.960 | .000075 | Middle Frontal | inferior frontal junction (IFJ) | L | -42, 13, 36 |
| 3.832 | .000127 |  | A8vl, ventrolateral area 8 | R | 42, 27, 39 |
| 3.457 | .000546 |  | A6vl, ventrolateral area 6 | L | -32, 4, 55 |
| 3.636 | .000125 |  | A10l, lateral area 10 | L | -26, 60, -6 |
| 4.505 | .000007 | Orbital | A12/47o, orbital area 12/47 | R | 40, 39, -14 |
| 3.718 | .000201 | Inferior Temporal | A20cl, caudolateral of area 20 | L | -59, -42, -16 |
| 3.942 | .000081 | Fusiform | A20rv, rostroventral area 20 | L | -33, -16, -32 |
| 3.427 | .000610 |  | A37mv, medioventral area 37 | L | -31, -64, -14 |
| 3.558 | .000373 | Parahippocampal | A35/36c, caudal area 35/36 | R | 26, -23, -27 |
| 3.582 | .000341 | Superior Parietal Lobule | A5l, lateral area 5 | R | 35, -42, 54 |
| 3.688 | .000226 | Inferior Parietal Lobule | A39c, caudal area 39(PGp) | L | -34, -80, 29 |
| 3.845 | .000120 |  | A39c, caudal area 39(PGp) | R | 45, -71, 20 |
| 3.991 | .000065 | Precuneus | A7m, medial area 7(PEp) | L | -5, -63, 51 |
| 4.701 | .000002 | Occipital Lobe | mOccG, middle occipital gyrus | L | -31, -89, 11 |
| 4.48 | .000007 |  | mOccG, middle occipital gyrus | R | 34, -86, 11 |
| 3.940 | .000081 |  | OPC, occipital polar cortex | R | 22, -97, 4 |
| 4.795 | .000002 |  | msOccG, medial superior occipital gyrus | L | -11, -88, 31 |

|  |  |  |  |  |  |  |
| --- | --- | --- | --- | --- | --- | --- |
|  | 3.488 | .000487 |  | msOccG, medial superior occipital gyrus | R | 16, -85, 34 |
|  | 3.615 | .000301 |  | lsOccG, lateral superior occipital gyrus | L | -22, -77, 36 |
|  | 3.36 | .000755 |  | lsOccG, lateral superior occipital gyrus | R | 29, -75, 36 |
| **SCR > Fear** |  |  |  |  |  |  |
|  | -3.355 | .000794 | Amygdala | Medial and lateral amygdala | R | -23, -3, -20 |
|  | -3.844 | .000121 | Inferior frontal | A44v, ventral area 44 | R | 54, 14, 11 |
|  | -4.068 | .000047 | Orbital | A11m, medial area 11 | L | -6, 52, -19 |
|  | -3.864 | .000112 | Paracentral Lobule | A4ll, area 4, (lower limb region) | L | -4, -23, 61 |
|  | -3.860 | .000113 | Superior Parietal Lobule | A7r, rostral area 7 | R | 19, -57, 65 |
|  | -3.551 | .000384 |  | A7c, caudal area 7 | R | 19, -69, 54 |
|  | -3.446 | .000569 | Postcentral | A1/2/3ulhf, area 1/2/3(upper limb, head and face region) | R | 50, -14, 44 |
|  | -3.441 | .000579 | Insular | G, hypergranular insula | L | -36, -20, 10 |
|  | -4.317 | .000016 |  |  | R | 37, -18, 8 |
|  | -4.598 | .000004 |  | dlg, dorsal granular insula | R | 39, -7, 8 |

Our results are in line with multiple previous findings indicating a positive relationship between the subjective fear ratings and autonomic responses ^16–19^. However, here we also showed that the brain regions involved in the accurate prediction of these two measures are possibly distinct. For instance, brain regions such as the amygdala, insula, and ventromedial prefrontal cortex appeared mostly involved in the prediction of physiological reactivity (Fig. 4b, right panel) while regions of the middle frontal gyrus, dorsomedial prefrontal cortex, and lateral orbital cortex were more closely related to the subjective reports of fear (Fig. 4b, left panel). This suggests that some caution may be warranted in the use of physiological reactivity as the sole source of information to infer the subjective suffering associated with fear and anxiety disorders.

Our results raise the question of the relation between physiological reactivity and subjective fear in the brain. To what extent are their representations independent? Is the subjective fear rating a late-stage readout of the physiological reactivity? Similar questions have been previously discussed in the consciousness literature ^20,21^. For instance, Maniscalco & Lau ^21^ tested multiple models formalizing the potential relations between sensory signal and subjective judgment. Their results suggest that a hierarchical model in which subjective experience depends on late-stage read-out best accounted for the data. This is notably in line with higher-order ^3,20,22^ and constructivist theories ^23,24^ of emotions suggesting that first-order representations (possibly reflected by the physiological reactivity) may need to be attended or meta-represented downstream for subjective experiences to occur. This is also in accord with a recent review of the literature ^25^ indicating that a meta-representation of the lower-level affective processes might be implemented by the middle frontal gyrus and other areas in the lateral prefrontal cortex.

Further evidence for a hierarchical model comes from results indicating that low-level affective processes do not seem necessary to generate a conscious experience of fear. For instance, patients presenting bilateral lesions of the amygdala have been reported to be capable of experiencing fear in some specific situations ^26,27^. Although there is still some debate regarding the possible mechanisms leading to these subjective experiences, the overall evidence at least does not seem incompatible with higher-order models.

A higher-order perspective is in line with previous results ^12,26–29^ and ours, but models of emotions are still being debated ^30^. For instance, one may argue against this hierarchical or higher-order view based on experimental demonstrations that the electrical stimulation of the amygdala itself can trigger a subjective experience of fear and anxiety ^31^. While this demonstration was compelling, it is important to mention that in that study this phenomenon occured only in 1 out of 9 patients. This inconsistency may be partly attributed to inter-individual differences in the spread of electrical activity to other brain regions. However, it is worth noting that the stimulation had a clear dose-dependent effect on the objective physiological response, which was observed across the entire group of patients. Taken together, these results suggest that the amygdala might play a central role in generating physiological responses but possibly a marginal role in generating the conscious experience of fear.

One reason for the skepticism about higher-order models might be that anxiety disorders have been reliably associated with a dysregulation of physiological reactivity ^32^. As such, higher-order structures are typically conceptualized as playing more of a complementary role in these pathologies. However, it is worth noting that the therapeutic success of psychotherapies for anxiety and depression appears to be mediated by brain regions such as the dorsomedial prefrontal cortex, posterior cingulate gyrus, precuneus and some regions of the temporal lobes ^33^. Also, recent findings indicated that the inhibition of the amygdala by the dorsolateral prefrontal cortex was positively associated with the outcome of exposure therapy ^34^. As such, some higher-order processes may also have an important incidence for therapeutic success.

One challenge in the implementation of a higher-order approach to anxiety disorders is the reliance on self-reported measures. Fear is indeed multifaceted and, as a result, ratings can be influenced by multiple factors such as arousal, proximity of threat, and other negative emotions. Furthermore, the means of fear assessment can greatly influence the outcome. We opted for offline categorical ratings as this is the typical approach for clinical diagnosis, but one may worry that this may not directly reflect the online subjective experience of fear. Our results suggest that both assessment methods may at least partly reflect similar processes as our decoders trained to predict offline ratings could predict weakly but significantly online ratings in an independent validation fMRI task (Supplementary Methods and Fig. S2). However, further work may be needed to determine precisely which aspects of fear are more salient with different means of assessment and how to cover accurately the multiple dimensions relevant to the self-report of fear.

Another concern is that emotional states have been proposed to involve (and sometime interfere with) cognitive functions ^35^. As such, we can expect part of our results to represent this interaction rather than a strict representation of fear *per se*. For instance, the middle frontal gyrus has also been involved in the cognitive regulation of emotion ^36,37^ and in the regulation of the physiological reactivity network ^34^. Furthermore, activity in this region has also been associated with working memory and the retrieval of semantic information ^38^. The same logic applies to attentional processes with reported influences in the occipital, frontal, parietal, and ventral temporal regions ^39,40^. Because our experiment involved cognitive functions such as working memory and attention, our results may be partly associated with the interference of fear with these cognitive functions ^41^. This observation does not undermine our claim as complex interactions between cognitive and affective processes might also represent an important mechanism of change in psychotherapy ^42^ that requires further empirical investigation.

Given that multivoxel decoding involves many parameter choices, one may wonder if our results robustly generalize or if they are due to idiosyncratic details. Overall, our impression is the main results do hold up under different analyses (see Fig. 3b, S1 and S2 as well as Supplementary Results). One specific concern is with respect to the choice of a between-or within-subject decoding strategy. Here, both approaches presented some similarities, at least regarding subjective ratings of fear (see Supplementary Methods, Results and Fig. S4). However, the generally weaker performance of within-subject decoders rendered a direct comparison impractical, especially for skin conductance reactivity which contains too few trials within each participant. Therefore, throughout we primarily focused on the between-subject decoding approach.

Another concern pertains to the use of either binned data or single-trial data. Binned (or averaged) data can be useful to train the decoders as the process of averaging can remove some of the within-subject noise and can make the data manageable for the training procedure. However, it also appears important to test the accuracy of the decoder in the prediction of raw single-trial data. This is why we chose to combine both approaches and to also test our decoders on raw single-trial data (Supplementary Fig. S1).

Another important concern pertains to the generalizability of decoders to other datasets. Our decoders presented good generalization to an independent validation dataset (see Fig. 3b) but also weaker performances on a dataset coming from a different fMRI task (Supplementary Fig. S2). Training the decoders using data from multiple tasks would potentially allow to build decoders that could generalize better across different tasks and datasets.

In sum, we have exploited an opportunity to directly compare how machine learning decoders can predict the subjective fear rating and its correlated physiological activity. Our results suggest that the study of fear and anxiety disorders may benefit from a greater inclusion of subjective measures as they might index higher-order processes not readily accessible when studying physiological reactivity alone. This may prove to be an important means to optimize treatments and further tailor interventions to specifically alleviate the subjective suffering associated with fear and anxiety disorders.

## Methods

### Participants

The discovery cohort included thirty-one participants (15 females; mean age = 23.29; SD = 4.21). Participants were included if they reported, on a 6-point Likert scale, “high” or “very high” fear of at least one animal included in the experiment (see Stimuli and Task for a detailed list). Skin conductance reactivity was not acquired for four participants and technical issues prevented from recording the skin conductance of two participants. As a result, the data of twenty-five participants were available to train the skin conductance reactivity decoder. The first independent validation cohort (same task) included twelve participants (2 females; mean age = 25.75; SD = 3.98) and skin conductance reactivity was acquired for 8 of them. The second independent validation cohort (different task) comprised 17 participants from the discovery cohort (5 females; mean age = 21.92; SD = 1.54) that performed a different experimental task (see Supplementary Methods). Skin conductance reactivity was recorder for all participants. All participants provided written informed consent and the study was approved by the Institutional Review Board of Advanced Telecommunications Research Institute International (ATR), Japan.

### Stimuli and Task

The experimental procedure has been described in full detail elsewhere ^43^. Briefly, participants underwent a 1-hour fMRI session where they were presented with images of the most commonly feared animals (e.g., snake, spider, cockroach, bee, bat, mouse, dog, cat, shark, etc.) as well as pictures of other animals and objects. We chose to present 90 different images per category and to include 30 animal categories and 10 object categories (for a total of 3,600 different images). The 30 different animal categories included reptiles (snake,turtle, and gecko), amphibians (frog), insects (cockroach, beetle, ant, spider, grasshopper, caterpillar, bee, butterfly, and fly), birds (robin, peacock, and chicken), annelids (earthworm), mammals (mouse, guinea pig, bat, dog, sheep, cat, rabbit, horse, and giraffe) and aquatic animals (shark, whale, common fish, and dolphin). The database also included 10 categories of human-made objects (airplane, car, bicycle, scissor, hammer, key, guitar, cellphone, umbrella, and chair). The images presented a full frontal view of the object or animal and no other recognizable object was clearly identifiable in the background. Images were cropped so that they would frame the object. The final images were 533 × 533 pixels and covered 13.33 degrees of visual angles during the procedure. The average contrast and luminance of images were not different between categories ^43^. The data of the human-made objects were not analyzed. Trials were organized in six runs of 600 trials interleaved with short breaks. The sequence of presentation was pseudo-randomized and fixed across participants.

### MRI parameters

Participants were scanned in two 3T MRI scanners (Prisma Siemens and Verio Siemens) with a head coil at the ATR Brain Activation Imaging Center. During the experiments, we obtained 33 contiguous slices (TR = 2000 ms, TE =30 ms, voxel size = 3 × 3 × 3.5 mm^3^, field-of-view = 192 × 192 mm, matrix size = 64 × 64, slice thickness = 3.5 mm, 0 mm slice gap, flip angle = 80 deg) oriented parallel to the AC-PC plane, which covered the entire brain. We also obtained T1-weighted MR images (MP-RAGE; 256 slices, TR = 2250 ms, TE = 3.06 ms, voxel size = 1 × 1 × 1 mm^3^, field-of-view= 256 × 256 mm, matrix size = 256 × 256, slice thickness = 1 mm, 0 mm slice gap, TI = 900 ms, flip angle = 9 deg.).

### Recording of electrodermal activity

Skin conductance reactivity was determined during the fMRI sessions using BrainAmp Ag/AgCl sintered MR electrodes (Brain Products). The electrodes were disposed on the distal phalanges of the index and middle fingers of the left hand. Skin conductance reactivity was determined in response to the first image of each chunks of images of a given category. Following previous methodologies ^43^, we determined the maximum amplitude in a time window of 1 to 5 seconds following the image onset and removed from this value the baseline activity in a 2-second window before the image onset. Responses smaller than 0.2 microsiemens (μS) were recoded as 0 (see Supplementary Methods). Responses were square-root transformed to correct for the skewness of the distribution ^44^. This standard analytical procedure allowed for our results to be readily put in correspondence with previous findings. However, this approach presents the disadvantage of allowing for the peak of skin conductance reactivity of one trial to occur within the time window of the following trial. We quantified that this scenario happened on 2.62% of all trials. This did not prevent us from developing a sensitive and accurate decoder of the skin conductance reactivity (see Fig. 3, 4, S1, and S2). However, this represents a source of noise that could be avoided in future experiments by using longer presentation chunks.

### Comparing subjective fear ratings and skin conductance reactivity

To determine the correlation between subjective fear reports and skin conductance reactivity, we first established, for each participant, an average level of skin conductance reactivity for each animal category. Since the first trial of each run (i.e., trials were organized in six runs of 600 trials) was typically associated with greater skin conductance reactivity, we removed these six trials as they didn’t represent typical reactivity to the image category *per se*. This removed six out of the 720 trials. The remaining trials were winsorized (5th and 95^th^ percentile), averaged within category and standardised. We then established group-level mean value for each category by averaging across participants. This was achieved both for the subjective fear ratings and for the skin conductance reactivity. The mean categorical values were then standardised at the group level. These standardised values were correlated to determine the association between subjective fear ratings and skin conductance reactivity at the group level. The results are presented in Figure 2.

### Preprocessing of fMRI data

The fMRI images captured during the experiment were realigned to the first fMRI image, coregistered, and motion-corrected (using 6 motion parameters) in SPM 12 (Statistical Parametric Mapping; www.fil.ion.ucl.ac.uk/spm) ^45^. Functions of pyMVPA (www.pymvpa.org) ^46,47^ implemented in the Neurodebian environment ^48^ were used to remove the linear trend and to deconvolve the signals using the least-square separate approach ^49,50^. This method allowed to iteratively fit a general linear model to estimate the brain response to the first presentation of each chunk of images. Each general linear model includes one parameter modeling the current trial, and two parameters modeling all other trials in the design. Via this method, we were able to obtain one parameter estimate (i.e., a beta image) for each individual trial of our rapid-event related design (720 beta images for each participant). Data were also normalized to the MNI space and smoothed (FWHM = [8,8,8]) using SPM 12.

### Developing whole-brain decoders

In order to build between-subject decoders, we aimed at creating binned beta images for each participants that would represent the levels of our outcomes (i.e., subjective fear ratings and skin conductance reactivity). This approach was also used in order to average out some of the between-trial noise. As such, to train the subjective fear decoders, the binned beta images were created by binning together (i.e. averaging), within-participant, the trials corresponding to the same level of fear (0= ‘No fear” to 5 = “Very high fear”). A similar procedure was used to create the binned beta images to train the skin conductance reactivity decoder. We aimed at creating six binned images per participants to reflect the different skin conductance reactivity levels. However, because of the skewness of the distribution, splitting the data according to even quintiles would result in an over representation of the trials with very small reactivity (i.e., most of the trials are below 0.2 microsiemens). As such, trials below 0.2 microsiemens were considered to be part of the binned beta image of level 0. The remaining trials were grouped into quintiles (computed individually) and the binned beta images 1 to 5 were obtained by averaging the corresponding images together. The number of trials in each bin was used to set the number of trials randomly selected to constitute the binned image of level 0. Binned beta images were mean centered within-subject.

We first trained whole-brain decoders using support vector regression in a leave-one-subject-out cross-validation procedure (implemented in Matlab [https://www.mathworks.com/products/matlab.html] using the CanlabCore toolbox [https://github.com/canlab/CanlabCore] and the Spider machine learning library [http://people.kyb.tuebingen.mpg.de/spider/main.html]). The predicted values of each beta image were used to establish the area under the ROC curve (AUC) of the decoders. To determine the statistical significance of the AUC, we conducted a permutation test by randomly permuting (1,000 times) the labels of the beta images in the datasets. Applying the decoders to this permuted data allowed to obtain a distribution of AUCs under the null hypothesis. This was achieved to obtain a critical value for significance at p = .05 (dashed lines in Fig. 3a and Fig. 4b).

### Testing whole-brain decoders on categorical beta images

If the whole-brain decoders can truly predict subjective fear ratings and skin conductance reactivity, their predictions of the different categorical images should correlate with the real observed values. In order to compare the decoding results with the behavioral data, we generated binned categorical images by removing the first trial of each block from the binned beta images (see above *Comparing subjective ratings and skin conductance reactivity*). This resulted in removing 6 trials out of the 720 beta images. We submitted the average categorical images to the whole-brain decoders. The predicted values were winsorized (5th and 95th percentile) and standardized within participant. At the group level, these values were averaged, standardized and then correlated with the mean subjective fear ratings and skin conductance reactivity (see Fig. 3c).

### Within-region decoding

The brain regions were defined using a parcellation of the cortex based on functional connectivity ^15^. We selected the 210 cortical regions of the atlas as well as the amygdala and hippocampus, for a total of 214 regions. We iteratively trained decoders within each of the selected regions to predict either one of the outcomes. This procedure provided us with a correlation coefficient between the predicted and real values for each decoder, within each region. This allowed for a direct comparison of the correlation coefficients between decoders using Fisher’s method ^51^. As such, this procedure was used to determine where in the brain one decoder presented a better performance than the other (e.g., a better prediction of the subjective ratings than the skin conductance reactivity). The false discovery rate of this series of dependent tests was controlled using the method described by Benjamini & Yekutieli ^52^ and implemented using the Matlab toolbox fdr_bh (http://kutaslab.ucsd.edu/matlabmk_fn_docs/matlabmk/fdr_bh.html). To facilitate the interpretation, we plot in Figure 4a the difference in the AUCs of both decoders within each region.

## Data availability

The data supporting the main findings of this study will be made available on the ATR website and from the corresponding authors upon reasonable request.

## Acknowledgements

The study was conducted in the ImPACT Program of Council for Science, Technology and Innovation (Cabinet Office, Government of Japan). This work was partially supported by ‘Brain machine Interface Development’ under the Strategic Research Program for Brain Sciences supported by the Japan Agency for Medical Research and Development (AMED). This study was also partly funded by the US National Institute of Neurological Disorders and Stroke of the National Institutes of Health (grant no. R01NS088628 to H.L.). V.T-D. is supported by a fellowship from the Fond de Recherche du Québec - Santé (FRQS) and by a fellowship from the Canadian Institutes of Health Research. We thank K. Nakamura and M. Miuccio for their help in scheduling and conducting the experiment, Dr K. Ide and Dr T. Chiba for help with the recruitment and the diagnosis of patients, N. Hiroe and H. Moriya for assistance with equipment, and Y. Shimada and A. Nishikido for operating the fMRI scanner. We also thank Yoni K. Ashar, Luke J. Chang and the participants of the Methods In Neuroscience at Dartmouth (MIND) summer school 2017 for helpful discussions regarding brain decoding and the use of the CanlabCore toolbox.

## Author Contributions

V.T-D., M.K and H.L., designed the study, V.T-D implemented the experiment and conducted the experiment, V.T-D. analysed the results and V.T-D., M.K., and H.L wrote the manuscript.

