## Supplemental Information for "Multivoxel Pattern Analysis Reveals Dissociations Between Subjective Fear and Its Physiological Correlates"

**Affiliations:**
1- Department of Decoded Neurofeedback, ATR Computational Neuroscience Laboratories, Kyoto, 619-0288, Japan.
2- Department of Psychology, UCLA, Los Angeles, 90095, USA.
3- Faculty of Information Science, Nara Institute of Science and Technology, Nara, 630-0192, Japan.
4- Brain Research Institute, UCLA, Los Angeles, 90095, USA.
5 - Department of Psychology, University of Hong Kong, Pokfulam Road, Hong Kong
6 - State Key Laboratory of Brain and Cognitive Sciences, University of Hong Kong, Hong Kong

**Table of Contents**

**Supplementary Methods** **3**
- Apparatus for stimuli presentation 3
- Testing whole-brain decoders on single-trial data 3
- Testing whole-brain decoders on an independent validation dataset 3
- Testing whole-brain decoders on specific phobia patients 4
- Assessing the performances of the whole-brain decoders on single-trial images 5
- Within-subject decoders 6

**Supplementary Results** **7**
- Testing whole-brain decoders on single-trial data 7
  - Area Under the Curve 7
  - Mixed effect models 7
- Testing whole-brain decoders on an independent validation dataset 7
  - Area Under the Curve 8
  - Mixed effect models 8
- Patients presenting specific phobias 8
- Within-subject decoders 9

**Supplementary Discussion** **9**

**Supplementary Figures** **11**
- Figure S1. Whole-brain decoders present sensitive and specific predictions of single-trial data. 11
- Figure S2. Whole-brain decoders present weaker, but statistically significant predictions of an independent validation dataset. 12
- Figure S3. Decoding accuracy of single-trial images in patients presenting specific phobia. 13
- Figure S4. Within-Subject decoding of the subjective fear ratings. 14

**Supplementary References** **15**

### Supplementary Methods

#### *Apparatus for stimuli presentation*

Visual stimuli were projected on a translucent screen by an LCD projector (DLA-G150CL, Victor). The projector spanned 20 × 15 deg in visual angle (800 × 600 resolution) and had a refresh rate of 60 Hz. The experiment was conducted using the PsychoPy2 software (v1.83) (1).

#### *Testing whole-brain decoders on single-trial data*

To determine the sensitivity of the whole-brain decoders to predict single-trial data, we used a similar leave-one-subject-out decoding approach, but here the test dataset was the single-trial data of the left-out participant. This procedure ensured that the data of the test participants were not included in the training dataset. The single-trial data were selected from the 720 trials that were the first of each series of images of a same category. To allow an unbiased assessment of the performance of the decoders using mixed effect models, the test datasets were balanced to avoid creating skewed distributions. For the subjective fear ratings, we sub selected, for each participant, an even number of random trials with low (< or equal to ratings of two) and high fear (> or equal to the rating of three). Regarding the skin conductance reactivity, we balanced the dataset using the quintile procedure described above, but applied the decoders to the unaveraged (e.g., single-trial) beta images (i.e., single-trial beta images).

#### *Testing whole-brain decoders on an independent validation dataset*

To determine how the whole-brain decoders could generalize to the independent validation data, we tested the prediction of the whole-brain decoders on data from another fMRI experiment. The experiment is described in detail elsewhere (2). Briefly, in two fMRI sessions, 17 participants were presented with a total of 40 images of “feared animals” and 20 images of “non feared animals” and objects. Offline categorical ratings obtained prior to the experiment were used to determine the animal categories to determine has “fearful” and “non fearful”. Animal categories associated with “High” or “Very High” fear were selected to be the fearful categories. Following the presentation of each image, fear ratings were collected in the scanner using a 7-point likert scale. Skin conductance was also recorded during the experiment and the reactivity to each image was established using the same procedure as described in the Methods section.

The preprocessing of the fMRI images was also similar to the preprocessing used in the main experiment: the images were realigned, coregistered, normalized to the MNI space, and smoothed (FWHM = [8,8,8]) using SPM 12. Since this design was not a rapid-event related design (i.e., images were presented for six seconds on the screen), we didn’t use the least-square separate approach and conducted first-level GLMs with one regressor modeling each trial and their time derivative. The resulting single-trial beta images were attributed either their subjective fear rating (i.e., online ratings) or their skin conductance reactivity level. The skin conductance was determined using the procedure described above (i.e., threshold of 0.2 microsiemens and individual quintiles). Beta images were then standardised and used to test the predictions of the whole-brain decoders.

***Testing whole-brain decoders on specific phobia patients***

One may expect anxiety disorders to be associated with significantly different brain processes then neurotypical participants (3). As such, one specific concern is whether the whole-brain decoders could predict accurately the brain data of patients presenting specific phobia. To tackle this question, we recruited three patients (3 females; Mean age = 26.33; SD = 7.77) who met the DSM-IV criteria for specific phobia for at least one animal in our database. Patients were diagnosed using the Structured Clinical Interview for DSM-IV conducted by three medical doctors trained in psychiatric assessment. Diagnoses were established by inter-rater agreement. The patients completed the same fMRI procedure as the neurotypical participants and were presented with 3600 images of animals and objects. As a result, some of the presented images were depicting the feared animal. Following the procedure described above, we obtained single-trial images of the patients. These images were submitted to the whole-brain decoders in order to determine their capacity to predict the subjective fear and skin conductance responses of patients.

***Assessing the performances of the whole-brain decoders on single-trial images***

For each participant and for each decoder, the predictions of the whole-brain decoders were assessed using the area under the ROC curve. This was established in the discrimination of low (< or equal to two) and high (> or equal to three) values of each outcome. The values of the area under the curve were submitted to a repeated-measure ANOVA with two within-subject factors (Decoders and Test dataset) and their interaction. Paired-sample t-tests were also carried out as follow-up analyses.

We also used a second approach to determine if the fear ratings and skin conductance reactivity (i.e., real values) could be accurately predicted by the decoders. For each decoder, we carried out a two-level mixed effect model predicting the real values of the outcome using three fixed effects (predicted values of the decoder, test dataset, predicted values X test datasets).We also allowed the intercepts to have a random component in order to correct for a potential clustering of error within-participants (4). The predicted values of the decoders were standardized within participant. This approach allowed to first test for an interaction between the predicted values and the test dataset in the prediction of the real values. Follow-up analyses were carried using two-level mixed effect models predicting the real values using one fixed effects (predicted values) and one random component (the within-participant intercept).

***Within-subject decoders***

Between-subject decoders allow to leverage data coming from a great amount of participants and can potentially generalise to new participants. However, between-subject decoders present the disadvantage of potentially missing some of the fine-grained brain representations as the data required for these analyses need to be smoothed significantly to facilitate between-subject correspondence. For this reason, we also trained within-subject decoders to determine if the fine-grained brain representations also followed a similar pattern as the one observed using the between-subject decoders. This however poses a challenge regarding the skin conductance reactivity decoder because a much smaller volume of data is available since skin conductance responses occurred on average on 28.21% of trials. As a result, we present here only the within-subject decoders of the subjective fear ratings and tried to determine if a similar pattern of accuracy could be observed as with the between-subject decoders. Accordingly, we conducted within-subject decoding in the significant regions previously reported in Figure 4b and tried to determine if similar results could be observed within-subject (i.e. better decoding of subjective fear ratings in the significant regions of the middle frontal gyrus than the significant regions of the amygdala, insula and ventral middle prefrontal cortex).

The data used to train within-subject decoders were preprocessed in the same way as the data used to train the between-subject decoders, but were not smoothed. Within-subject data were binned as a function of block and subjective fear ratings. We used support vector regression and a leave-one-block-out cross-validation approach.

### Supplementary Results

#### ***Testing whole-brain decoders on single-trial data***

*Area Under the Curve*

The results of the repeated-measure ANOVA showed a significant decoder X test dataset interaction ($F(1,24) = 62.17$; $P < .0001$; two-sided) (see Figure S1). Each decoder presented greater areas under the curve when tested on their corresponding dataset (Subjective fear decoder: $t(24) = 4.68$; $P < .0001$; two-sided; Skin conductance reactivity decoder: $t(24) = -7.55$; $P < .0001$; two-sided) (see Figure S1 a and b). The effect sizes were considered to be of large sizes both for the subjective fear decoder (Cohen's $d = 0.94$) and for the skin conductance reactivity decoder (Cohen's $d = 1.51$) (corrected for dependence between the means) (5).

*Mixed effect models*

Likewise, we observed a significant interaction between the predicted values and the test datasets both for the subjective fear (predicted values X test datasets interaction: $t(16757) = -8.01$; $P < .0001$; two-sided) and the skin conductance reactivity decoders (predicted values X test datasets interaction: $t(14720) = -14.12$; $P < .0001$; two-sided). These results indicate that both decoders present a better prediction of their corresponding dataset at the single-trial level. More precisely, the subjective fear decoder predicts more accurately the real values of the subjective fear dataset ($t(11681) = 23.77$; $P < .0001$; two-sided) than the skin conductance dataset ($t(5076) = 5.01$; $P < .0001$; two-sided). Similarly, the skin conductance reactivity decoder predicts more accurately the real values of the skin conductance reactivity dataset ($t(5076) = 20.17$; $P < .0001$; two-sided) than the subjective fear dataset ($t(9644) = 4.46$; $P < .0001$; two-sided).

***Testing whole-brain decoders on an independent validation dataset***

*Area Under the Curve*

The results of the repeated-measure ANOVA showed no significant decoder X test dataset interaction ($F(1,16) = .669$; $P = .426$; two-sided). There was also no statistically significant difference between the test datasets for each decoder (Subjective fear decoder: $t(17) = .775$; $P = .450$; two-sided; Skin conductance reactivity decoder: $t(17) = -612$; $P = .549$; two-sided). The mean values were although qualitatively in the same direction as the ones obtained with the single-trial data (see Figure S2).

*Mixed effect models*

Likewise, we observed no interaction between the predicted values and the test datasets both for the subjective fear (predicted values X test datasets interaction: $t(1306) = -1.03$; $P = .30$) and the skin conductance reactivity decoders (predicted values X test datasets interaction: $t(1306) = -1.59$; $P = 0.11$; two-sided). However, both decoders present a statistically significant prediction of their corresponding dataset. More specifically, the subjective fear decoder presented a statistically significant prediction of the subjective fear dataset ($t(803) = 2.87$; $P = .004$; two-sided) and not of the skin conductance dataset ($t(503) = 1.35$; $P = .17$; two-sided). Similarly, the skin conductance reactivity decoder present a statistically significant prediction of the real values of the skin conductance dataset ($t(503) = 2.38$; $P = .018$; two-sided) and not of the subjective fear dataset ($t(805) = -0.019$; $P = .98$; two-sided) (see Figure S3 c and d).

***Patients presenting specific phobias***

Whole-brain decoders were used to predict the single-trial images of the patients. The area under the curves of the predicted values are presented in Figure S3. To quantify the divergence of patients with respect to the group of participants, we used the standard deviation of the distributions. For the subjective fear decoder, the area under the curve were included within 0.53 and -0.5 standard deviations for both test datasets. With respect to the skin conductance reactivity decoder, the area under the curves of patients are included within 1.85 and -0.56 standard deviations for both test dataset. While this is a restricted sample, these results suggest that whole-brain decoders do not perform significantly worse when predicting data of patients. The results even suggest that the skin conductance reactivity of some patients might be predicted more efficiently than the rest of the group (z scores of 1.38 and 1.85).

***Within-subject decoders***

Within-subject decoders were trained to predict the subjective fear ratings in the brain regions previously reported as significant in Figure 4b. The results are presented in Figure S4. A similar tendency as with between-subject decoders can be observed: the significant regions of the middle frontal gyrus tend to present a greater accuracy in the prediction of the fear ratings than the other significant regions of the amygdala, insula and ventromedial prefrontal cortex ($t(30) = 1.79$; $P = .08$; two-sided). It is also to be noted that the within-subject decoders were significantly less accurate than the between-subject decoders ($t(8) = 2.35$; $P = 0.046$; two-sided). This is to be expected as within-subject decoders are trained with a subsample of the data.

### Supplementary Discussion

The supplementary analyses provided information regarding the accuracy of whole-brain decoders to predict the single-trial images both of the discovery cohort and the independent validation. The results indicate that whole-brain decoders present sensitive and specific predictions of the single-trial data of the discovery cohort. Also, when whole-brain decoders are applied to the independent validation data, their predictions are weaker, but both decoder still present a statistically significant prediction of their corresponding dataset. This weaker performance is to be expected since the independent validation dataset was acquired using a different fMRI task that potentially engaged different brain processes. As such, during this experiment, participants were not required to perform a 1-back task on image category and were explicitly asked to rate their subjective fear of the presented images. Also, the independent validation dataset was substantially smaller than the original dataset which likely decreased the statistical power of these tests.

The supplementary analyses also suggested that the whole-brain decoders might present similar levels of accuracy in patients presenting specific phobia. More precisely, no patients presented prediction accuracies significantly below the group mean (all above -.53 standard deviation). While these results were obtained with a small number of patients (N = 3) they still provide some evidence suggesting that the whole-brain decoders of subjective fear ratings and skin conductance reactivity might generalize to patients. Although, further analyses will be necessary to assess this claim with greater statistical rigour.

Lastly, the supplementary analyses indicate a tendency for a better decoding of subjective fear ratings from the middle prefrontal cortex as opposed to the amygdala, insula, and ventromedial prefrontal. Although, the interpretation of this marginally significant result is limited by the relative weakness of the decoding performance in the within-subject analyses. This weak decoding performance and the small number of skin conductance responses obtained within-subjects prevented further investigation of the difference in accuracies of the within-subject decoders. As a result, further investigations will be needed to provide more conclusive evidence regarding the role of the fine-grained information in the decoding of subjective fear ratings and skin conductance reactivity.

### Supplementary Figures

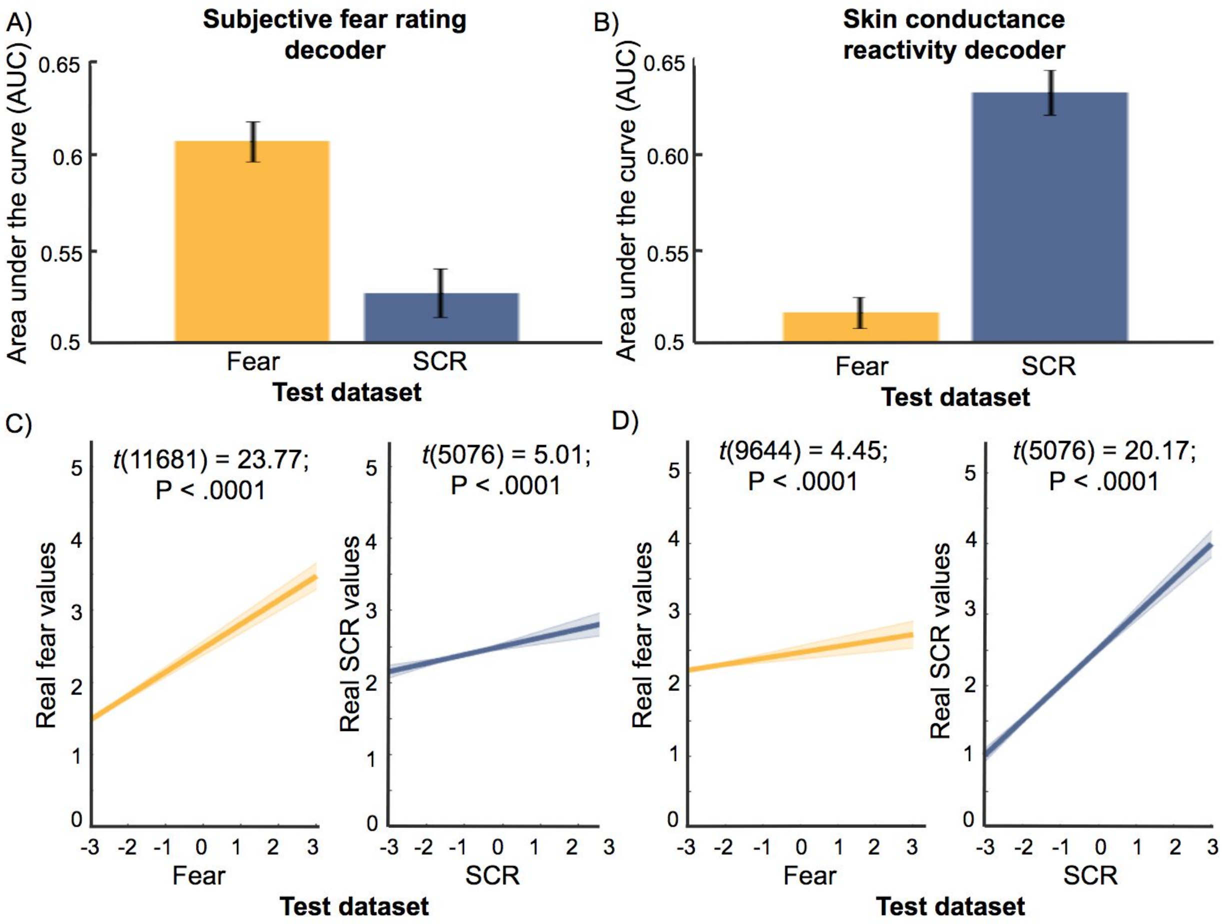


***Figure S1. Whole-brain decoders present sensitive and specific predictions of single-trial data.***

Whole-brain decoders of the subjective fear rating (left panels) and the skin conductance reactivity (right panels) present more accurate predictions when tested on the dataset they were trained to predict (e.g., subjective fear decoder predicting fear data). This is illustrated both by the use of area under the curves (top panels) and by mixed-effect models (bottom panels). Error bars are +/- 1 S.E.M. Shaded error bars correspond to the 95% confidence intervals of the slope and intercept of the mixed effect models.

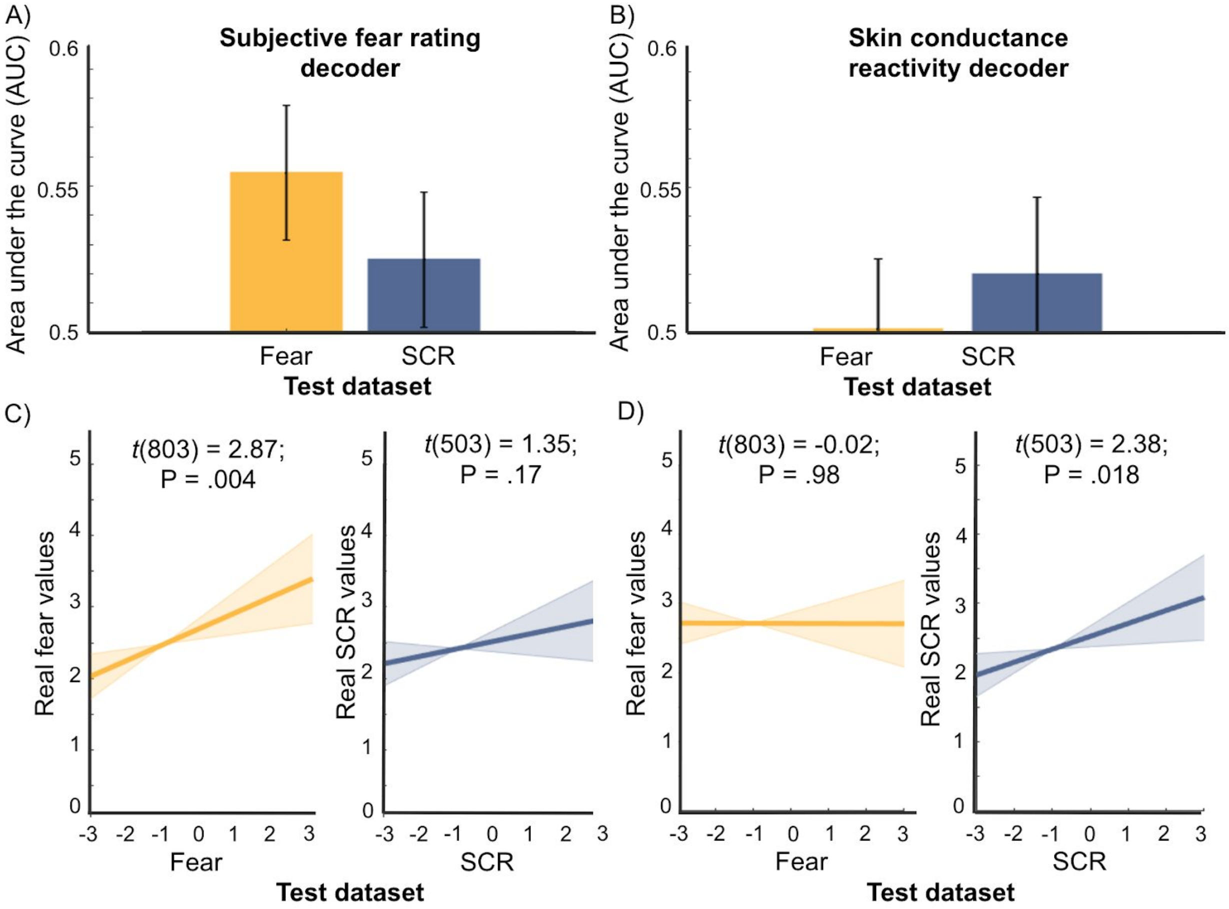


***Figure S2. Whole-brain decoders present weaker, but statistically significant predictions of an independent validation dataset.***

Using the area under the ROC curves (a and b), the whole-brain decoders of the subjective fear rating (left panels) and the skin conductance reactivity right panels) do not present a significant prediction of their corresponding dataset. Although, mixed-effect models (c and d) indicate a weak, but statistically significant capacity of the decoders to predict their corresponding datasets. These results are qualitatively in line with the single-trial results of the discovery cohort (i.e., greater capacity to predict the corresponding dataset). Error bars are +/- 1 S.E.M. Shaded error bars correspond to the 95% confidence intervals of the slope and intercept of the mixed effect models.

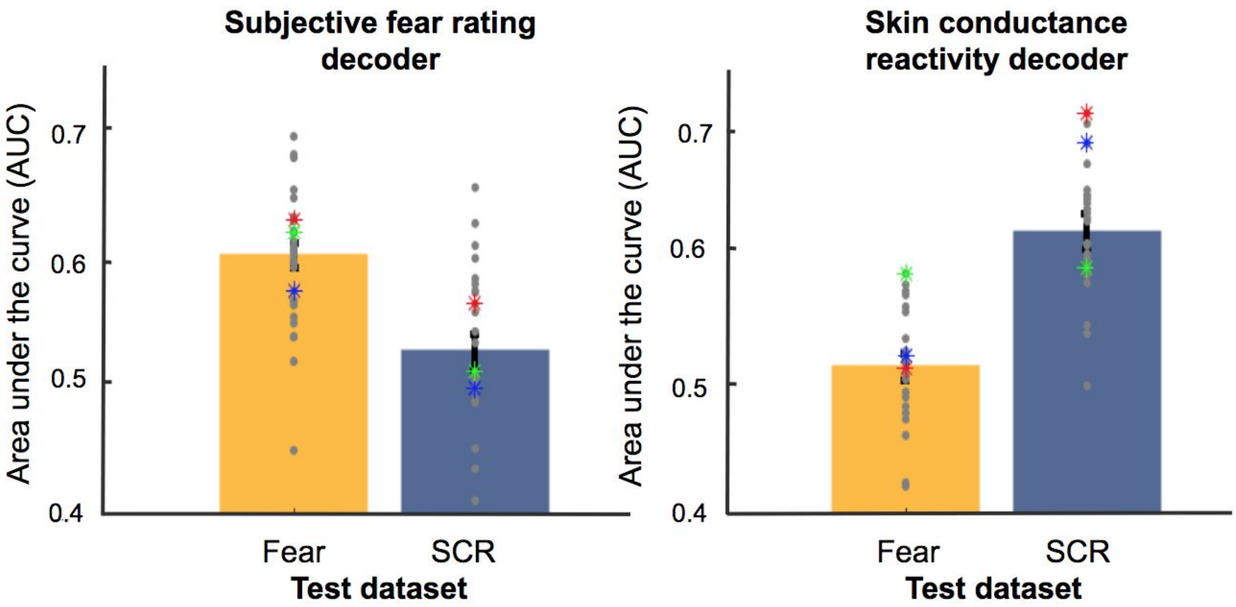


***Figure S3. Decoding accuracy of single-trial images in patients presenting specific phobia.***

Colored stars represent the area under the curve of patients presenting specific phobias. No patients presented area under the curves below 0.56 standard deviation. As such, these results do not indicate a clear decrease in the capacity to predict the single-trial values of patients. Error bars are +/- 1 S.E.M.

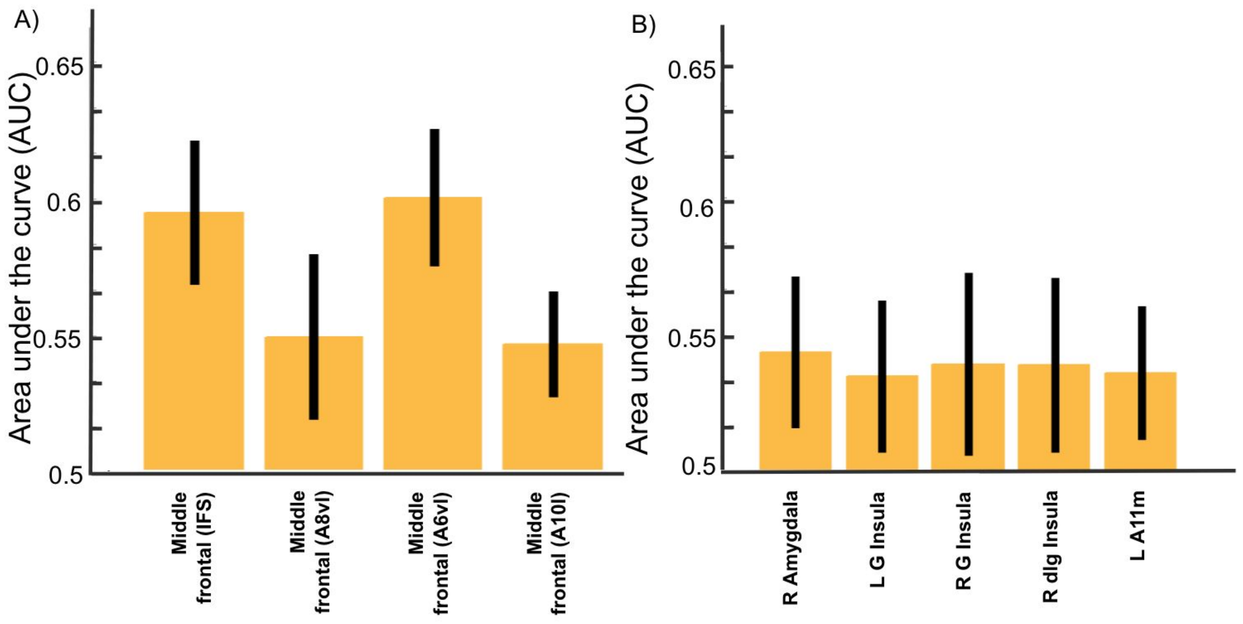


***Figure S4. Within-Subject decoding of the subjective fear ratings.***

Within-subject decoders were less accurate than the between subject decoders in the prediction of the subjective fear ratings ($t(8) = 2.35$; $P = 0.046$; two-sided). However, they still show a similar tendency to present better performances in the middle frontal gyrus than the amygdala, insula, and ventromedial prefrontal cortex ($t(30) = 1.79$; $P = .08$; two-sided). Error bars are +/- 1 S.E.M.
